# MetaMutationalSigs: Comparison of mutational signature refitting results made easy

**DOI:** 10.1101/2021.04.04.438398

**Authors:** Palash Pandey, Sanjeevani Arora, Gail Rosen

## Abstract

**Summary:** The analysis of mutational signatures is becoming increasingly common in cancer genetics, with emerging implications in cancer evolution, classification, treatment decision and prognosis. Recently, several packages have been developed for mutational signature analysis, with each using different methodology and yielding significantly different results. Because of the nontrivial differences in tools’ refitting results, researchers may desire to survey and compare the available tools, in order to objectively evaluate the results for their specific research question, such as which mutational signatures are prevalent in different cancer types. There is a need for a software that can aggregate results from different refitting packages and present them in a user-friendly way to facilitate effective comparison of mutational signatures.

**Availability and implementation:** MetaMutationalSigs is implemented using R and python and is available for installation using Docker and available at: https://github.com/EESI/MetaMutationalSigs

**Contact:** Gail Rosen.

**Supplementary information:** More information about the package including test data and results are available at https://github.com/EESI/MetaMutationalSigs

## Introduction

Mutational signature analysis provides an operative framework to understand the somatic evolution of cancer from normal tissue (Robinson et al 2020; Alexandrov et al, 2020; Brunner et al., 2019; Yoshida et al., 2020; Moore et al.,2020). From the earliest phases of neoplastic changes, cells may acquire several types of mutations in the form of single nucleotide variants, insertions and deletions, copy number changes and chromosomal aberrations. These mutations are caused by multiple mutational processes operative in cancer leaving behind specific footprints in the DNA that can by captured by mutational signature analysis(Alexandrov et al.,2013; Alexandrov et al, 2020). It is becoming increasingly evident that these mutational signatures are not only important for understanding cancer evolution but also may have therapeutic implications, thus this a very active and important area of research (Iqbal et al., 2020; Campbell et al., 2017; Chung et al., 2020; Alexandrov et al., 2020).

The basic idea behind mutational signatures is that mutational processes create specific patterns of mutations. Thus, it follows that if one can identify these patterns in a given sample then they can essentially detect the corresponding mutational processes. The possible mutations are grouped into 6 mutation types based on the base where the mutation was observed. These 6 mutation types are C>A, C>G, C>T, T>A, T>C, and T>G. Now, these 6 types of mutations are further divided based on their location, i.e., other bases that are in their immediate proximity providing the 96 mutation types that are termed the single base substitution (SBS) context. Alexandrov et al. first developed and applied this idea to The Cancer Genome Atlas (TCGA) data and identified the first iteration of 30 SBS signatures which were compiled into the Catalogue Of Somatic Mutations In Cancer (COSMIC) and came to be used as the de facto reference for signature refitting, we refer to these v2 signatures as COSMIC legacy SBS signatures (Campbell et al., 2020; Forbes et al., 2017). The initial study was then expanded to the analysis of data from the Pan-Cancer Analysis of Whole Genomes (PCAWG) project (Campbell et al., 2020), resulting in two additional signature classes with multiple signatures in each class. These new classes are COSMIC V3 SBS, double base substitutions (DBS) signatures and insertions/deletion (ID) signatures, which are in (Alexandrov et al., 2018).

The mutational signature analysis workflow involves multiple steps that require different amounts of time and processing power. Briefly, the workflow starts from Binary Alignment Format (BAM) files that are aligned to a reference genome and then proceeds to the variant calling step which outputs the Variant Calling Format (VCF) files. These steps are usually very resource-intensive and thus do not allow for much experimentation on personal computers; the downstream steps of variant filtering and annotation are much faster. The final step, the mutational signature analysis, is the least resource-intensive and, therefore, is easier for users to compare multiple methods on their desktop. Therefore, to facilitate comprehensive mutational signature refitting analyses, we developed the package, MetaMutationalSigs. We developed a wrapper for 4 typically used refitting packages (Rosenthal et al., 2016; Blokzijil et. al. 2018; Gori et. al., 2018; Wang et al., 2020), that have various underlying methodologies, such as Bayesian inference, non-negative least squares, and quadratic programming. Here, we have developed a standard format for inputs and outputs for easy interoperability and effective comparison, respectively. With our previous experience in visualization of genomic data (Lan et al., 2014), we have implemented standard visualizations for the results of all mutational signature packages to ensure easy analysis. MetaMutationalSigs software is easy to install and use through Docker.

### Approach

The two major methods typically used for mutational signature analysis are signature refitting and de-novo signature extraction. Signature refitting methods try to reconstruct the observed mutational pattern in the sample (the frequencies of 96 types of mutations) using linear combinations of known signatures (COSMIC Legacy SBS and COSMIC V3 SBS, ID, DBS, etc.), these methods work quite well on small sample sizes (such as single samples) and are widely used with small datasets (Omichessan et al., 2019). Signature extraction methods infer signatures from a given dataset, and then compare the extracted signatures with known reference signatures. Each extracted signature is assigned to a known signature if their cosine similarity exceeds a set threshold, otherwise signatures with similarity less than the threshold are ignored (Alexandrov et al.,2013). There are a few important caveats to signature extraction as recently discussed in (Omichessan et al., 2019): 1) a novel signature can be very similar to several reference signatures and the assignment is not always perfect and 2) the threshold for assignment plays a crucial role but is not widely agreed upon and using a different threshold can change the assignment (Omichessan et al., 2019).

We chose signature refitting as our primary task and implemented high performing packages as identified in (Omichessan et al., 2019) that were implemented in R using common input matrix generated using SigProfilerMatrixGenerator (Bergstrom et al., 2019). We implement DeconstructSigs (Rosenthal et al., 2016), MutationalPatterns (Blokzijil et. al. 2018), Sigfit (Gori et. al., 2018), Sigminer (Wang et al., 2020), these tools build up on other tools such as (Mayakonda et al., 2018; Huang et al., 2018). Our package outputs several data files in comma separated values (CSV) format ready for further analysis and visualization using external packages along with visualizations of the signature contributions as described in **Table 1**.

**Table 1.** Summary of result files.

| File Name | Format | Description |
| --- | --- | --- |
| Heatmap_contributions_all_sigs_legacy.pdf | pdf | Contributions for all COSMIC Legacy SBS signatures to the overall signature. |
| Heatmap_contributions_all_sigs_SBS.pdf | pdf | Contributions for all COSMIC V3 SBS signatures. |
| Heatmap_COSMIC_legacy.pdf | pdf | Heatmap for difference between the predicted contributions by different tools. One for each sample. |
| Heatmap_COSMIC_V3.pdf | pdf | Heatmap for difference between the predicted contributions by different tools. One for each sample. |
| legacy_pie_charts.html | html | Interactive pie charts of COSMIC legacy SBS contribution, per sample and for each tool. |
| sbs_pie_charts.html | html | Interactive pie charts of COSMIC V3 SBS signature contributions, per sample and for each tool. |
| legacy_rmse_bar_plot.pdf | pdf | Reconstruction error using COSMIC Legacy SBS signatures for each tool. |
| sbs_rmse_bar_plot.png | pdf | Reconstruction error using COSMIC V3 SBS signatures for each tool. |
| toolname_results\legacy_sample_error.csv | csv | Data used to create the bar plot. |
| toolname_results\legacy_sample_contribution.csv | csv | Data used to create heatmap and pie chart. |
| toolname_results\sbs_sample_error.csv | csv | Data used to create the bar plot. |
| toolname_results\sbs_sample_contribution.csv | csv | Data used to create heatmap and pie chart. |

We compare packages using the root mean squared error (RMSE) between the reconstructed and actual signals. RMSE is a performance metric commonly used in signal processing (Rosen, 2007).

## Discussion

The massive increase in the number of software packages has made managing dependencies quite burdensome, coupled with incompatible data formats for signature matrices can make mutational signature refitting results difficult and hard to compare. Our package, MetaMutationalSigs, provides a simplified approach for performing the setup related tasks so that more focus can be placed on the analysis. Investigators should keep in mind that refitting approaches need *a priori* knowledge about the samples and each package for effective interpretation (Maura et al., 2019), and the results should not be used as-is without an assessment of the cell biology and genomics.

Future work for this project would focus on expanding the tool to work with more packages and keep the reference signatures updated as new versions are released. Due to the open-source nature of the project, we also welcome additional feature requests using the project link on GitHub https://github.com/EESI/MetaMutationalSigs

## Ethics Statement

Not applicable to this study.

## Data Availability Statement

All test data used is open source and is available with the software at GitHub https://github.com/EESI/MetaMutationalSigs

---

**Figure 1.**
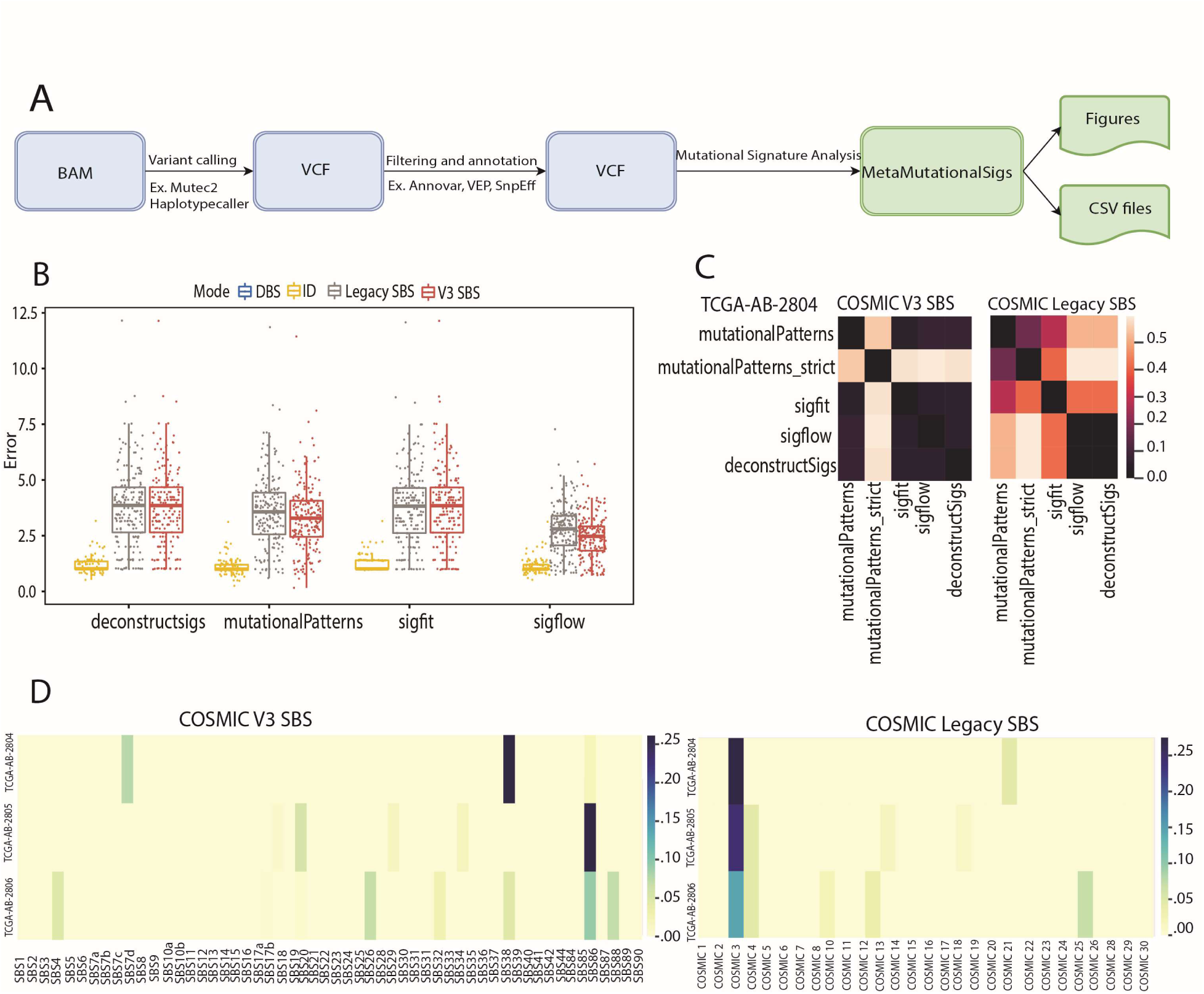
Workflow and results for metaMutationalSigs. **A)** The workflow for mutational signature analysis, starting with a BAM file of a sequenced genome or exome and followed by variant calling, filtering, and annotation. Our tool, MetaMutationalSigs, analyzes signatures found in a VCF. **B**) RMSE for each tool and reference signatures (lower values are better) for SBS and IDs (no tool predicted DBS for the samples used). While RMSE does not change for deconstructSigs and Sigfit, the RMSE significantly drops for mutationalPatterns and Sigflow with COSMIC v3. **C**) Heatmap of Euclidean distance between the predicted contributions of COSMIC v3 SBS vs. COSMIC legacy SBS signatures by different tools for the same TCGA patient sample. With the legacy signatures, tools are generally less in agreement in their resulting signature contributions, while with COSMIC v3 signatures, the standard use tools are all in agreement with each other. Sigflow had the lowest RMSE and was selected for analysis in **D. D**) Heatmaps of COSMIC v3 SBS vs. COSMIC legacy SBS mutational signature contributions using whole-exome sequence data from three TCGA patients with acute myeloid leukemia. Here, each row is a patient sample. <u>Left</u>. COSMIC v3 SBS refitting provides different dominant signature contributions, TCGA-AB-2804: unknown etiology, TCGA-AB-2805 and TCGA-AB-2806: unknown chemotherapy and different DNA mismatch repair signatures, SBS20 and 26, respectively. COSMIC Legacy SBS refitting provides signature 3 (failure of double-strand break-repair by homologous recombination) as the dominant signature for all samples. The COSMIC v3 SBS refitting reveals multiple mutational processes may be playing a role in the overall signature contribution than is found with the COSMIC Legacy SBS refitting. =

